# Honey bee queen health is unaffected by contact exposure to pesticides commonly found in beeswax

**DOI:** 10.1101/2021.04.24.441288

**Authors:** Alison McAfee, Joseph P Milone, Bradley Metz, Erin McDermott, Leonard J Foster, David R Tarpy

## Abstract

Honey bee queen health is crucial for colony health and productivity, and pesticides have been previously associated with queen loss and premature supersedure. Prior research has investigated the effects of indirect pesticide exposure on queens via workers, as well as direct effects on queens during development. However, as adults, queens are in constant contact with wax as they walk on comb and lay eggs; therefore, direct pesticide contact with adult queens is a relevant but seldom investigated exposure route. Here, we conducted laboratory and field experiments to investigate the impacts of topical pesticide exposure on adult queens. We tested dose-response relationships of six pesticides commonly found in wax: coumaphos, tau-fluvalinate, atrazine, 2,4-DMPF, chlorpyriphos, chlorothalonil, and a cocktail of all six, each dosed up to 32 times the concentrations typically found in wax. We found no effect of any treatment on queen mass or sperm viability. Furthermore, none of the 1,568 proteins quantified in the queens’ fat bodies (a major site of detoxification enzyme production) were differentially expressed. In a field trial with N = 30 queens exposed to either a handling control, a solvent control, or a pesticide cocktail, we again found no impact on queen egg-laying pattern, mass, or emergence mass of daughter workers. Further, of the 3,127 proteins identified in fluid from the spermatheca (sperm storage organ), none were differentially expressed. These experiments consistently show that at realistic exposure levels, pesticides commonly found in wax have no direct impact on queen performance, reproduction, or quality metrics. We suggest that previously reported associations between high levels of pesticide residues in wax and queen failure are most likely driven by indirect effects of worker exposure (either through wax or other hive products) on queen care or queen perception.

## Introduction

Queens are normally the sole reproductive female within a honey bee colony and the reproductive status of a queen directly influences the colony’s overall health; however, queen quality can be compromised by environmental stressors [1, 2]. A queen can lay up to two thousand eggs/day in the spring [3], and a queen’s egg-laying rate directly influences the number of worker offspring produced in a colony. Furthermore, due to their haplodiploid sex determination system, the production of viable honey bee workers requires the fertilization of eggs [3], thus queens invest substantial metabolic resources to maintain viable spermatozoa within their spermathecae [4–6]. Honey bee queens take part in nuptial flights soon after their emergence as adults and store the spermatozoa (sperm) they acquire for the duration of their lives [3]. Once exhausted of viable sperm, queens can no longer produce female worker offspring and are replaced by the colony [4]. Colony losses are consistently attributed to poor queens by beekeepers [7], and pesticide exposure is an environmental factor that is linked to declines in reproductive health and longevity [1, 8–14].

While fulfilling agricultural pollination services, honey bees are often exposed to agrichemicals, including pesticides [10, 15–17]. Moreover, the transport of commercial colonies to multiple farms within a single pollination season can further increase the potential for contact with pesticides [10], as pesticide exposure risk varies among different land uses [17, 18]. Foragers collect and store residue-containing food (nectar and pollen) inside the hive; therefore, pesticides present in the ambient landscape are therefore commonly found as chemical residues in beeswax and food resources [10, 17, 19]. Additionally, miticides applied directly into the nest as a control measure against the parasitic Varroa mite (V. destructor) are found as chemical residues in wax [10, 17]. While both beekeeper-applied miticides and agrichemicals are detected in hive matrices from commercial colonies, miticides tend to be the most dominant residues [10]. In both cases, however, the effects of in-hive exposures to these residues—individually and in combination—on honey bee reproductive health is an understudied aspect honey bee toxicology [20]. While residues in food resources can result in oral exposure, chemicals present in wax can be delivered through contact with the cuticle. Beeswax is the primary nest substrate on which bees live and store food, and beeswax is largely comprised of complex esters and fatty acids, produced by the workers glandular secretions [21]. The composition of beeswax is conducive for accumulating lipophilic molecules, including some pesticides [22, 23]. As a result, beeswax from commercial honey bee colonies may contain a large number of chemical residues, some at high concentrations. A large-scale, multi-pesticide residue screening survey (n=108) found an average of 10 different pesticide residues in a given sample, with miticides (coumaphos, tau-fluvalinate, and 2,4-DMPF - an amitraz degradation product) and insecticides (fipronil, deltamethrin, fenpropathrin, and permethrin) among the most hazardous [10].

The likelihood for a chemical to result in harm (risk) depends on both its exposure and toxicity. The Hazard Quotient (HQ) is a risk estimation approach for multi-pesticide risk, calculated by dividing the exposure (amount) of a pesticide by its respective toxicity (LD_50_), then summing the HQs for each pesticide in a mixture to estimate the cumulative hazard [24]. The approach does not account for pesticide interactions, as in most cases these are not well-defined. It has been previously reported that colonies containing wax with higher HQs had a higher incidence of queen events, with queenright colonies having an average HQ around 1,500 and colonies exhibiting the loss of a queen having an HQ of nearly 3,500 [10]. High exposures to these miticides in beeswax during development have been shown to influence queen health, including effects on pupal weight, sperm count, and sperm viability [8, 25]. For example, topical treatment of queens with imidacloprid, a neonicotinoid insecticide, has been previously shown to reduce stored sperm viability [1]; however, this is not one of the most common pesticide residues found within hives [10, 17] especially in wax because it is hydrophilic. Aside from mating flights early in life, queens typically remain within a colony, but they may still be exposed to in-hive pesticide residues from food (through the workers) and wax (the nest substrate). Indeed, some evidence suggests that topical exposure could affect queen sperm health and brood offspring survival [1, 26]. Here we used previously reported in-hive concentrations of pesticides found in beeswax in order to examine the risks these residues may pose to adult queen reproductive health in both laboratory and field conditions.

## Results

A potential route of pesticide exposure for queen honey bees is topical absorption through contact with residue-containing wax while walking and laying eggs. While topical exposure of workers has been traditionally employed when screening for honey bee pesticide toxicity (See Tier I assessment, USEPA 2014) [27], with few exceptions [1, 26, 28, 29], the effects of contact exposures on individual queens are seldom investigated. In order to test if topical exposures of different pesticides commonly found in wax affected queen quality metrics, we exposed queens to varying doses of six pesticides (coumaphos, fluvalinate, 2,4-DMPF, chlorothalonil, chloropyrifos, and atrazine) as well as a complete cocktail, then measured queen mass, sperm viability, as well as other morphometrics associated with reproductive quality (head width, thorax width, spermatheca width, and sperm counts). The doses ranged from 1x to 32x, where x is the median wax concentration for that compound as reported by Traynor et al. [10] (Table 1). Control queens were exposed to an equal volume of solvent control (acetone).

**Table 1:** Applications of each pesticide in terms of hazard quotients (HQ) and parts per billion (ppb). Each treatment is expressed as a multiplication of the corresponding amounts quantified from commercially surveyed wax [10].

| Pesticide | LD<sub>50</sub> | T1 (1x) |  | T2 (4x) |  | T3 (8x) |  | T4 (16x) |  | T5 (32x) |  |
| --- | --- | --- | --- | --- | --- | --- | --- | --- | --- | --- | --- |
|  |  | ppb | HQ | ppb | HQ | ppb | HQ | ppb | HQ | ppb | HQ |
| Fluvalinate | 4.32 | 4,310 | 998 | 17,240 | 3,991 | 34,480 | 7,981 | 68,960 | 15,963 | 137,920 | 31,926 |
| Coumaphos | 5.93 | 943 | 159 | 3,772 | 636 | 7,544 | 1,272 | 15,088 | 2,544 | 30,176 | 5,089 |
| 2,4-DMPF | 75 | 304 | 4.1 | 1,216 | 16.2 | 2,432 | 32.4 | 4,864 | 64.9 | 9,728 | 129.7 |
| Chlorothalonil | 111 | 361 | 3.3 | 1,444 | 13.0 | 2,888 | 26.0 | 5,776 | 52.0 | 11,552 | 104.1 |
| Chlorpyrifos | 0.08 | 2.7 | 35.4 | 10.8 | 141.7 | 21.6 | 283.5 | 43.2 | 566.9 | 86.4 | 1,133.9 |
| Atrazine | 98.5 | 5.4 | 0.05 | 21.6 | 0.22 | 43.2 | 0.44 | 86.4 | 0.88 | 172.8 | 1.75 |
| Cocktail | - | - | 1,200 | - | 4,798 | - | 9,596 | - | 19,192 | - | 38,384 |

A total of 168 queens were analyzed, although four queens perished during the experiment for reasons not associated with a specific pesticide or dose (**Supplementary Table S1**). We first checked that the fertility and morphometric measurements of these queens was consistent with previous research (**Figure 1a**). As expected, queen weight was significantly and positively correlated with head and thorax width, as well as spermathecal diameter (all Spearman correlations, p < α = 0.003 with Bonferroni correction), as previously demonstrated [30, 31]. Moreover, sperm viability and sperm count were also positively correlated (ρ = 0.507, p = 5.1 x 10^-12^). Next, we tested if pesticide treatment affected queen mass or sperm viability, as these are quality metrics which have been previously been shown to change in response to abiotic stressors [1, 2, 6]. We found no dose-response relationships of any of the compounds in relation to queen weight or sperm viability after correcting for multiple hypothesis testing (**Figure 1b & 1c; Table 2**).

**Figure 1.**
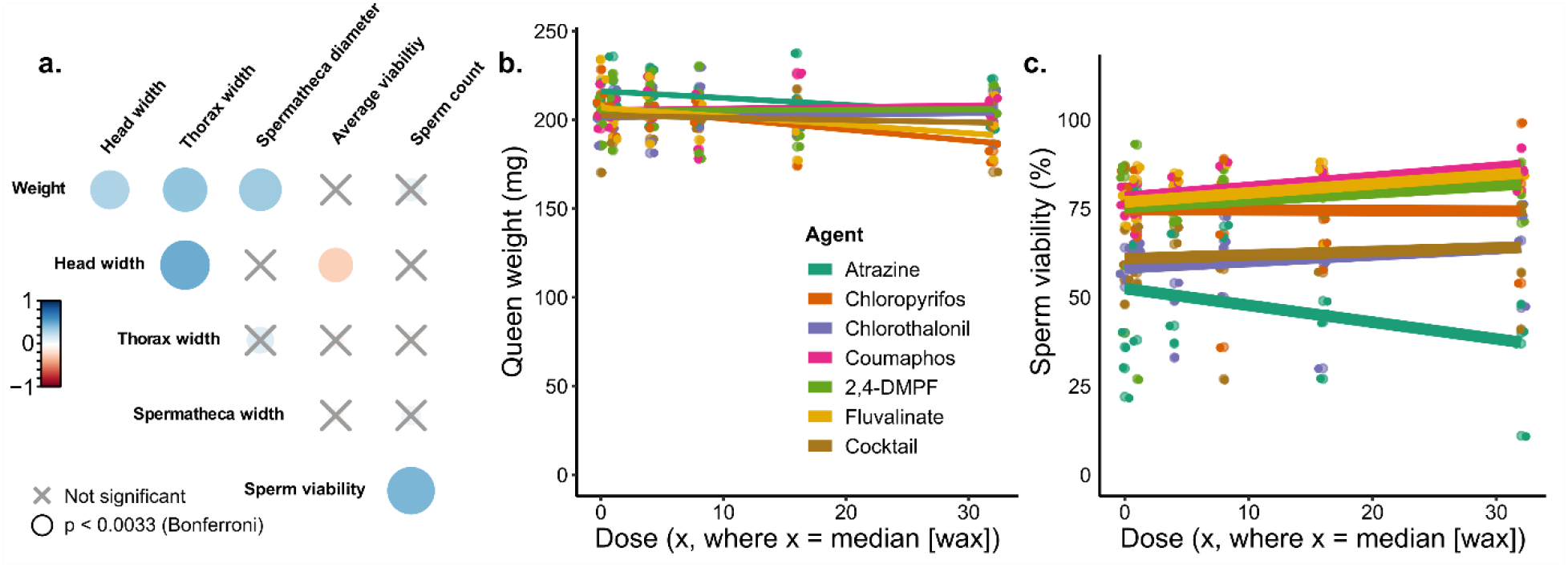
Evaluating the effect of pesticide doses on queen quality metrics. N = 4 queens were exposed topically (2 μl to the thorax, acetone solvent) to each pesticide and each dose, for a total of 168 queens. a) Queen quality metrics were recorded after exposure. The colour bar is proportional to the Pearson correlation coefficients. Neither queen weight (b) nor sperm viability (c) depends on dose or pesticide. See **Table 2** for a complete statistical summary.

**Table 2.** Summary statistics for dose-response queen exposures*

| Agent | R<sup>2</sup> | F | df<sub>1</sub> | df<sub>2</sub> | P value |
| --- | --- | --- | --- | --- | --- |
| Chlorpyrifos | 0.0232 | 1.18 | 3 | 20 | 0.342 |
| 2,4-DMPF | -0.0335 | 0.751 | 3 | 20 | 0.534 |
| Fluvalinate | -0.0454 | 0.696 | 3 | 18 | 0.567 |
| Coumaphos | 0.294 | 4.05 | 3 | 19 | 0.022 |
| Chlorthalonil | -0.119 | 0.218 | 3 | 19 | 0.882 |
| Atrazine | 0.116 | 2.01 | 3 | 20 | 0.146 |
| Mixture | -0.142 | 0.0878 | 3 | 19 | 0.966 |

\*Bonferroni correction: $p < \alpha = 0.007$ for significance

We hypothesized that queens would initiate a broad metabolic detoxification response following topical pesticide exposure. To test this hypothesis, we conducted mass spectrometry-based proteomics on the fat body tissue, a major source of detoxification enzyme production [32], of queens exposed to atrazine, coumaphos, and the cocktail treatment. We chose these specific treatments for analysis because, in addition to being a controversial endocrine disruptor [33, 34], atrazine is the major herbicide residue found in honey bee hive matrices [35] and may affect invertebrate reproductive characteristics [36]. Additionally, coumaphos was the compound most strongly (although still not significantly) correlating with sperm viability, and the cocktail treatment offers the most realistic exposure scenario. We quantified 1,568 protein groups in the fat body (1% protein and peptide FDR; **Supplementary Table S2 and S3)**, yet we observed no significant correlations between protein expression and pesticide identity or dose (analysis conducted using the limma package in R, see **Supplementary File S1** for example R code) even at a relatively loose false discovery rate (10%, Benjamini-Hochberg correction; **Figure 2a-d**).

**Figure 2.**
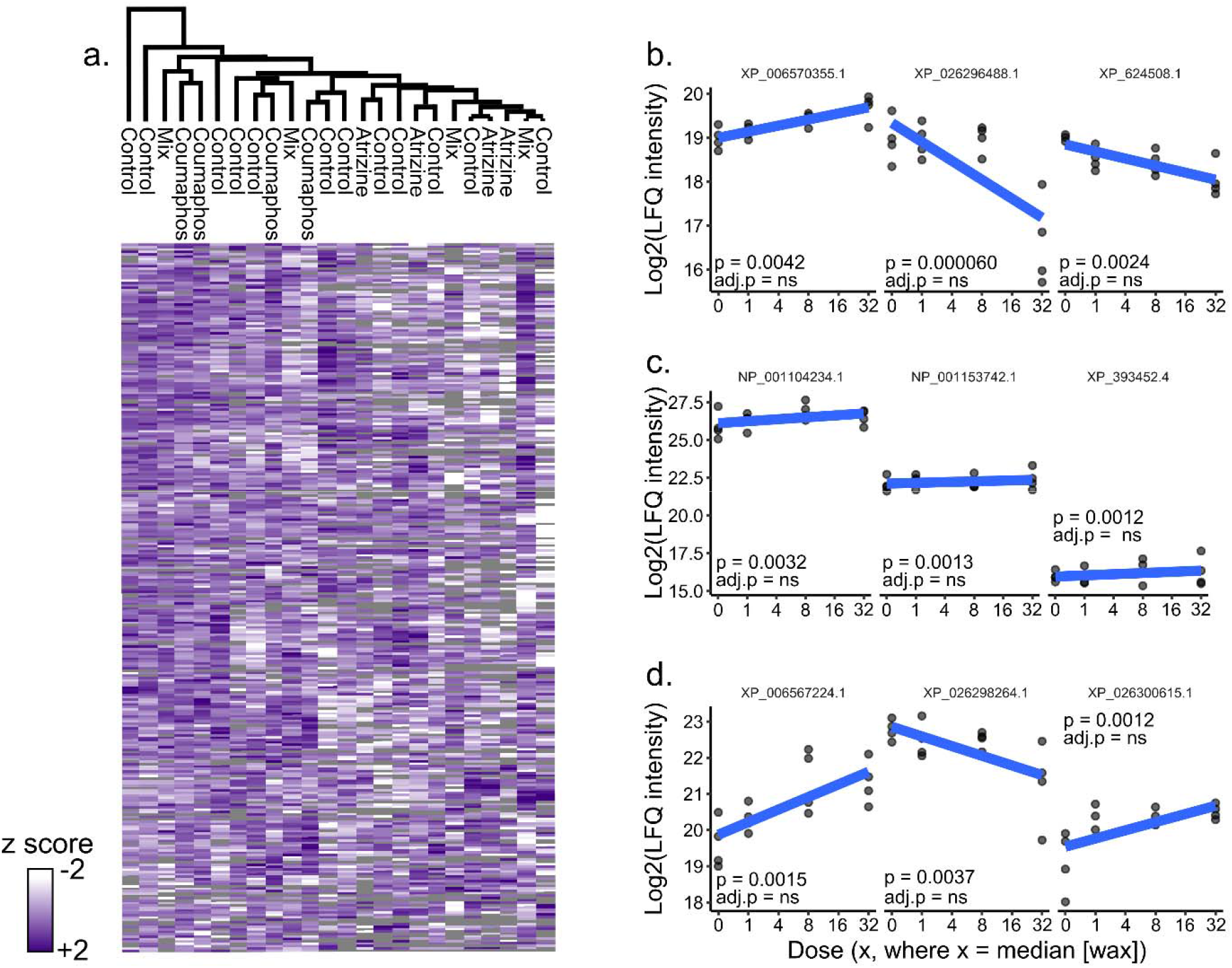
Evaluating the effect of pesticide exposure on queen fat body protein expression. We performed label-free quantitative (LFQ) proteomics on proteins extracted from fat bodies of atrazine-, coumaphos-, and cocktail-treated queens dosed at 0x, 1x, 8x and 32x. a) We found no differences between the highest pesticide dose (32x) [10] and their acetone controls (10% Benjamini-Hochberg correction). Samples and proteins were clustered via Euclidian distance, 300 clusters, 10 iterations. Grey tiles indicate samples in which a protein was not identified. Top proteins linked to pesticide dose for atrazine (b), coumaphos (c), and the cocktail (d). None was significant after correcting for multiple hypothesis testing. Accessions indicate refseq IDs. ns = protein does not survive 10% FDR.

Queen mass and sperm viability are important fertility metrics, but some of the impacts of pesticide on queen fertility can have a delayed response and require longer to manifest [11]. Moreover, some aspects of reproduction (e.g., laying pattern or vertical effects on progeny) are not captured by a laboratory dose-response experiment; therefore, we conducted a field experiment to measure additional queen and colony phenotypes before and after exposure to the pesticide mixture described above, but with modifications. Since we aimed to test only the cocktail and were thus not limited by additional queens needed to test each component individually, we were able to increase the complexity of the cocktail by the addition of fenpropathrin (insecticide), pendimethalin (herbicide), and azoxystrobin (fungicide)—three additional wax pesticide residues which were previously detected in >20% of beeswax samples [10]. We administered the cocktail using the same method as described for the dose-response experiment but delivered the cocktail at a Hazard Quotient (HQ) of ~3,500, which is the hazard level that has been previously found to be associated with ‘queen events’ (queen loss or supersedure) (**Table 3**) [10]. This corresponds to 2.3x of the median wax HQ found in [10]. During the experiment, four of the thirty original queens perished (unrelated to treatment group; one cocktail, two solvent, and one untreated queen). We measured the queens’ egg laying pattern and wet weight before and after pesticide treatment (**Supplementary Table S4**), as well as the average wet weight of newly emerged workers born from eggs laid before and after the queens were stressed (**Supplementary Table S5**). We analyzed these data as ratios (post-treatment:pre-treatment) to account for individual variation between the queens’ baseline characteristics, and we found no differences in any of these quality or performance metrics between the cocktail-treated queens and the solvent-treated controls or the untreated controls (linear model; sample sizes, F statistics, degrees of freedom, and p values are shown on the figures; **Figure 3a-c**). We further confirmed that the post-stress quality metrics were in the expected range; that is, the majority of queens had laying patterns near 100% coverage within the measured patch (80% or lower is considered to be a ‘poor brood pattern’ [37], which applied to only one of our queens), all but two queens had masses > 200 mg (within the range of previously published data [38–40]), and average callow worker mass was 107 mg (again, similar to previously reported data; [41]; **Figure 3d-f**).

**Table 3.**
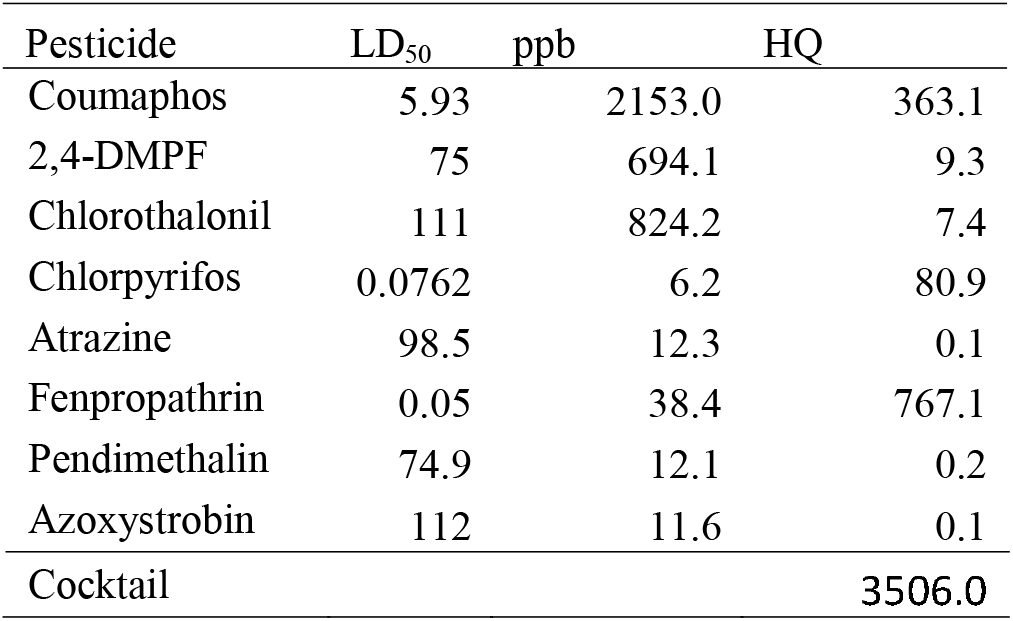
Composition of the 9-component cocktail used for the field trial, derived from [10].

**Figure 3.**
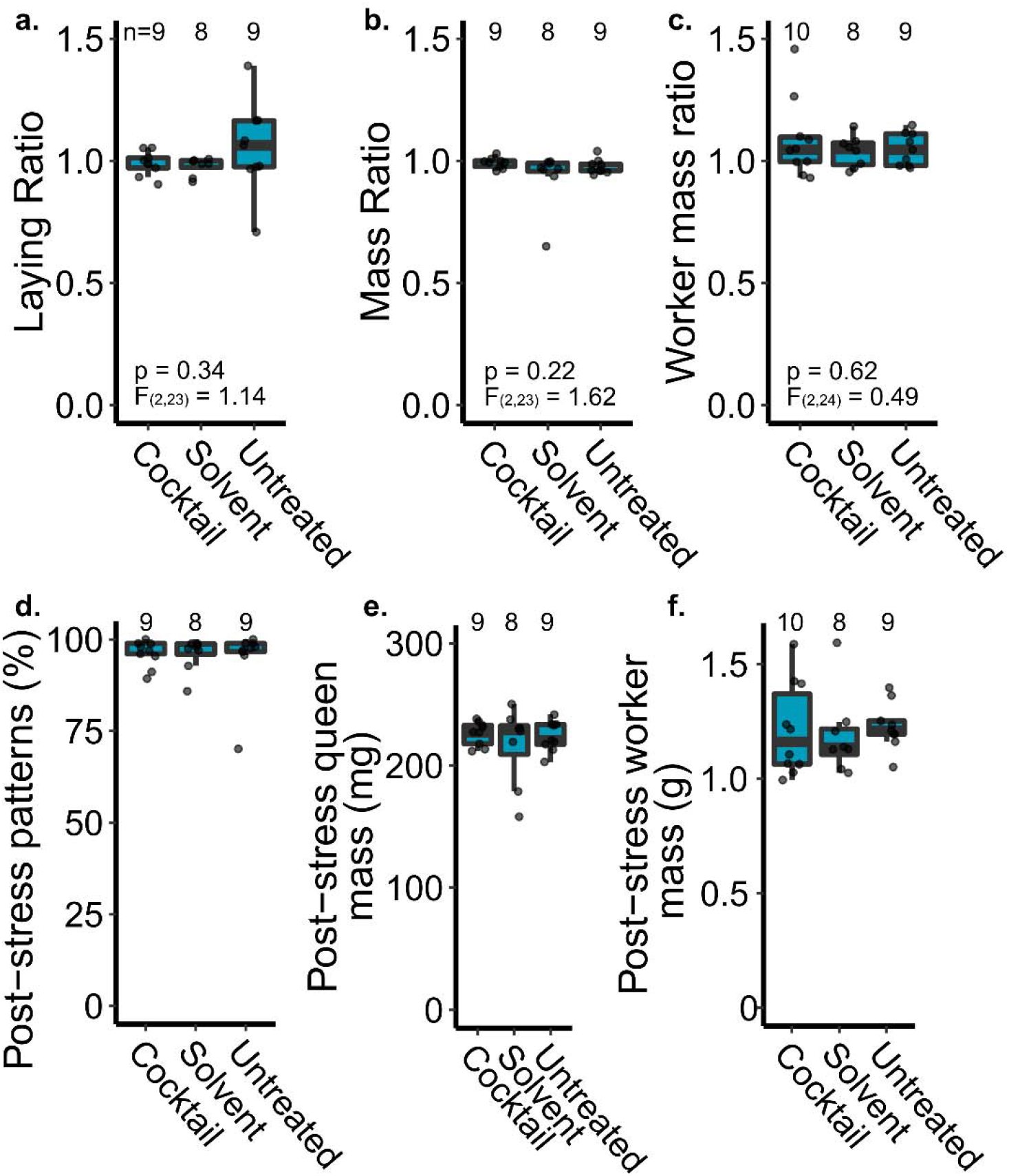
A field experiment evaluating the effect of topical pesticide cocktail treatment on queen performance. (a-c) We measured the queen’s laying pattern and mass immediately before and 2 weeks after topical pesticide exposure (2 μl, dosed at ~3,500 HQ, or 2.3x where x = the median wax concentration). Ratios indicate the post-stress metric relative to the pre-stress measurement to account for variation between individual queens. Sample sizes and statistical parameters are indicated on the graphs. (d-f) Non-normalized post-stress metrics. Boxes represent the interquartile range, bars indicate the median, and whiskers span 1.5 times the interquartile range.

We previously proposed candidate pesticide stress biomarkers expressed in the spermathecal fluid after topical pesticide exposure [42]: Catalase (XP_026296889.1) and Cytochrome c oxidase (XP_392368.1). We ultimately aimed to use these candidate markers to detect pesticide stress, among other stressors, in failed queens. Here, we evaluated proteins expressed in the spermathecal fluid in order to determine if these candidate biomarkers remain present in the expected expression pattern (elevated in pesticide-stressed queens) despite the longer post-stress recovery period relative to our previous experiment in which they were identified (2 weeks vs. 2 days) [42]. We identified 3,127 protein groups (1% protein and peptide FDR; **Supplementary Table S6**) but found no differentially expressed proteins in the spermathecal fluid of queens treated with the pesticide cocktail compared to the untreated or solvent-treated controls (**Figure 4a;** 10% Benjamini-Hochberg FDR). We also examined expression patterns of the candidate stress biomarkers specifically, but these too were not differentially expressed among groups (linear model, **Figure 4b & c)**, precluding their realistic utility as a diagnostic tool.

**Figure 4.**
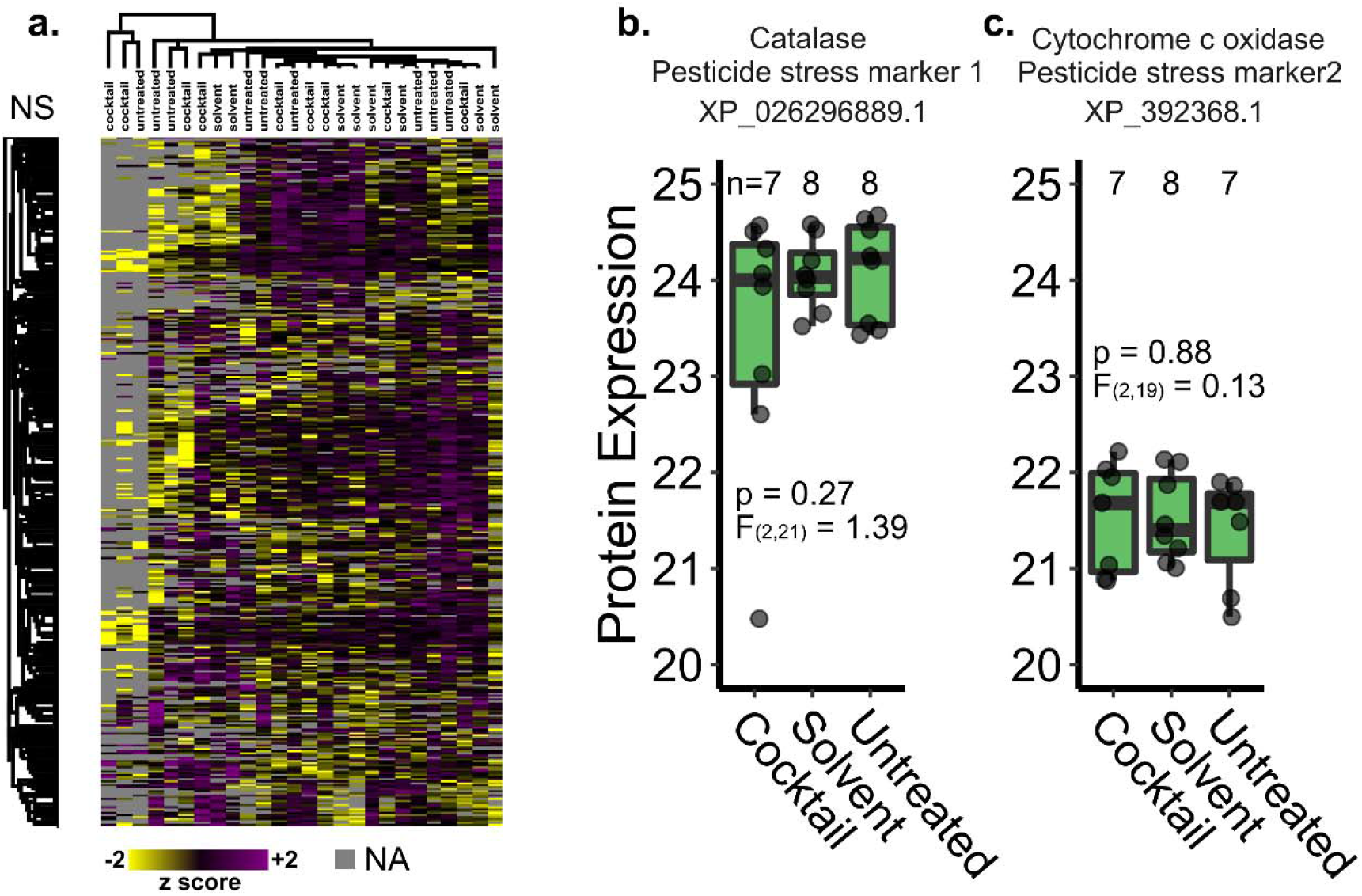
Protein expression in spermathecal fluid of queens treated with a pesticide cocktail. Queens were reintroduced to colonies after topical treatment (2 μl, dosed at 3,500 HQ) and sacrificed for analysis after 2 weeks. No proteins were differentially expressed at a global scale (limma, 10% FDR) (a) and the previously proposed pesticide stress biomarkers were also not differentially expressed (linear model; b & c). Samples and proteins clustered via Euclidian distance, 300 clusters, 10 iterations. Boxes represent the interquartile range, bars indicate the median, and whiskers span 1.5 times the interquartile range.

## Discussion

We report that adult queen health and sperm viability are likely unaffected by most pesticide residues commonly encountered through contact with beeswax. We report no substance-related effects from any of the six tested pesticides or their mixture on queen morphometrics, stored sperm quantity, or sperm viability at any dose levels (**Figure 1**). The herbicide atrazine caused the largest reduction in overall sperm viability across tested compounds, but even this effect was not significant (Table 2).

Interestingly, the treatment mixture of all pesticides combined at their highest dose-level resulted in lower queen sperm viability (mean: 62.0% SD: ±16.3%), which was lower relative to all other individual pesticides except for the herbicide atrazine (mean: 34.1% SD: ±15.9%), although this relationship was also not significant (Table 2). We observed no interactions (*i.e.*, additivity, synergy, or potentiation) among pesticides in our cocktail treatment. While it is not clear exactly how queens are so tolerant to topical pesticide exposure, these data suggest that direct pesticide exposure through contact with contaminated wax is not likely to adversely impact queens.

The highest exposure examined in this study was 32 times the respective median detection within beeswax from commercial colonies for each respective chemical (Table 1) [10]. These test concentrations exceeded maximum detections for all chemicals except chlorothalonil and 2,4,-DMPF, and the combination of all chemicals at these concentrations did not impart a measurable effect on tested queens. The exposures used in this experiment are also likely to be far above the theoretical pesticide exposures faced by an individual queen from contact from contaminated beeswax inside of a colony, because it can be assumed that only a portion of the residues present within beeswax are freely transferable to a queen’s cuticle. However, the dynamics of in-hive pesticide movement and how they shape colony exposure remain largely understudied, including how sub-lethal chronic exposure may affect a queen over the course of her lifetime.

Our laboratory results identifying no effect of topical exposure on queen quality metrics were corroborated by our field observations, where we again did not identify any impact of pesticide cocktail exposure on queen performance or average mass of her adult progeny (**Figure 3**). Furthermore, we did not observe exposure-associated ‘queen events’ in the weeks following reintroduction to their hives. Although it is possible that our observation period (2 weeks post-stress) was not long enough to observe changes in performance, we think this is unlikely because topical exposure of a neonicotinoid pesticide to queens has been previously shown to reduce sperm viability within 1 week [1], and other abiotic stressors can affect queen quality within days [2, 6, 43]. Together, these data suggest that topical pesticide exposure, such as what may occur from contact with wax, is not likely to be an important exposure route for queens.

The honey bee queen invests considerable metabolic resources in maintaining viable sperm within the spermatheca, and stored sperm is an important indicator of honey bee queen longevity [4–6, 44]. Our findings indicate that adult queens may have a robust capacity for protecting sperm from the potentially deleterious impacts of high concentration contact exposure to certain pesticides commonly found within commercial colonies. Another study, using similar experimental methods, also found no effects on sperm viability at exposures up to 100x the median detection for the miticide coumaphos [1]. These findings highlight that queen losses previously associated with residues in beeswax [10] are likely not the result of direct toxic effects on adult queen sperm health and instead may arise from other indirect social effects. One explanation may be that workers inhabiting hives that contain highly contaminated beeswax or pollen may perceive their environment in a way that makes them more likely to initiate the processes which result in queen events (queen replacement and death). Contrary to many other animals that exhibit increased cooperation under stressful conditions, it has been previously shown that workers that were starved during larval development emerge as adults with a reduced response to queen mandibular pheromone [45]. Queen pheromones are an important signal for modulating cooperative behaviors in workers such as foraging and brood rearing [3], and stressor-mediated changes to pheromone signaling deserve future attention. Oral pesticide exposure has been shown to influence queen nutrition during development [46], and it is unknown how pesticide exposure may interact with the queen pheromone production and their perception by workers. Finally, the most visible phenotypes under investigation as indicators of queen health (*e.g.*, brood pattern and colony population) are not solely under queen control, and it has been found that colony brood viability may be impacted by other stressors not relating to queen quality [47].

We previously observed that at a lower dose of the same pesticide cocktail (HQ 512), many proteins were differentially expressed in the queen’s spermathecal fluid within 2 days after exposure [42]. Furthermore, research by Chaimanee *et al*. identified acute detoxification responses as little as 1 day after exposure to imidacloprid, coumaphos, and amitraz [1, 28]. Here, however, we found that 2 weeks after exposure to a much higher dose (HQ 3,500) there were no discernable expression differences (**Figure 4**). This suggests that, at least for the compounds we tested here, queens may have a rapid detoxification response and quickly clear the compounds before harmful effects are realized. Unfortunately, we found that the protein biomarkers for pesticide exposure that we previously proposed do not have sufficient longevity to indicate exposure in a realistic scenario, and given that we found no direct effect of topical exposures on queen quality. A different strategy will be needed to 1) understand the basis for the relationship between high residue concentrations and queen events, 2) develop an exposure method that reflects that relationship, and 3) analyze queens exposed using those methods to suggest new candidate biomarkers.

## Conclusion

Our laboratory exposure data and field observations suggest that contact exposure of the pesticides most commonly found in wax is unlikely to directly impair queen quality. While we acknowledge that results of chronic exposure over the course of a queen’s lifetime may differ, we tested a range of doses far exceeding what a queen should ever encounter, and still we found no effect on queen quality metrics and fat body protein expression. Combined with observing no change in queen performance within colonies after exposure to the complete pesticide cocktail, these findings suggest that previous associations between residue concentrations and ‘queen events’ are more likely to be driven by indirect effects on the queen through exposed workers.

## Methods

### Queen dose-response exposures and dissections

Queens were purchased from Wilbanks apiaries and banked for approximately 2 weeks prior to the start of the experiment. Queens were placed in lots of 24 queens, of which four queens were exposed to one of six treatment doses consisting of a control (acetone only), a logarithmic ascending dose of the focal compound (six different compounds were tested), or cocktail of all chemicals. The dosing of these compounds was informed from residue data reported by Traynor *et al*. [10]. The six compounds we tested were the same as those used in the topical exposures for the field trial, except that we excluded fenpropathrin, azoxystrobin, and pendimethalin to simplify our experimental design. Our dosing ranged from 1x (where x = the median wax concentration) up to 32x (see **Table 1** for all doses). The cocktail was mixed such that each compound was present in the same relative concentration as reported by Traynor *et al*. [10] and at the same dosing scale as the individual compounds.

Two microliters of pesticide solution diluted in acetone were applied to the thorax of each queen, which were observed for approximately 5 minutes and then stored in Benton cages with attendants and water delivered by a damp dental wick in an incubation chamber kept at 34.5 °C. Queens were monitored and water replenished periodically during this time, approximately every 24 hours. 48 hours following exposure, queens were anesthetized with CO_2_ from sublimated dry ice until immobile. Each queen was removed from its cage, weighed, pinned to a dissection stage ventral-side up, and photographed. Then the abdominal tegument was separated at the 6th abdominal tergite and the spermatheca was gently removed, which was itself photographed prior to immersion in 1 mL of Buffer D (17⍰mM D-glucose, 54⍰mM KCI, 25⍰mM NaHCO_3_, 83⍰mM Na_3_C_6_H_5_O_7_). The queen was then stored in a 1.7 mL centrifuge tube at −80 °C for future fat body proteomics analyses.

### Sperm viability analysis

The spermatheca was ruptured with a pair of ridged forceps and the released contents mixed, transferred into a 1.7 ml centrifuge vial, and vortexed gently for 15 s. An aliquot of 200 μl was transferred to a 1.0 ml amber chromatography vial containing 2.0 μl propidium iodide solution and Sybr 14 dye from a Thermo-Fisher Live-Dead Sperm viability kit essentially as previously described [48]. This aliquot was vortexed to mix and capped, allowing it to incubate at room temperature until all queens were processed, which took approximately 2.5 h.

After the spermathecal contents had incubated with the dyes for at least 10 min and no more than 3 h, 20 μl was transferred to a Nexcelom Cellometer counting chamber for spermatozoa count and viability imaging essentially as previously described [11, 48]. The average of the three reads was taken for the measures of total spermatozoa concentration, and the viability was recorded as the ratio of live to total spermatozoa.

### Morphometric analysis

Photographs of queen morphometric measurements were analyzed using ImageJ The width of the head was taken at the widest point perpendicular to the body axis in a line passing over the frons. The width of the thorax was taken as the width of the mesothoracic sternite directly parallel to a line intersecting the tegulae. Measurements were converted from pixels to mm by comparison to a 0.1 mm microscopic rule.

### Pesticide stress field trial

Honey bee colonies were established from imported Tasmanian packages as part of a previously described experiment [49]. Briefly, packages were installed in standard 10 frame deep hive bodies, supplied with pollen and syrup, and after 1.5 months (two brood cycles), we split the colonies into 30 three-frame nucleus colonies (nucs), each with one frame of honey, one frame with open brood, and one frame with capped brood. We supplied each nuc with a frame feeder for light syrup (~35% sucrose) and a ½ lb pollen patty (15% protein), which we fed continuously throughout the duration of the field trial. All nucs were kept in a single apiary in Richmond, Canada.

Each nuc was supplied with a queen imported from California (all from a single shipment). Caged queens were placed between two frames with the cage screen facing down and allowed to acclimate for 3 days, at which time the queens were released. 2 days later, we checked the nucs for queen acceptance and those that were rejected were supplied a new queen from the same batch.

Once each queen had been laying eggs for at least 2 weeks, we evaluated their laying patterns, as previously described [49], by locating a patch of approximately 100 eggs and recording how many cells within that patch were apparently missed (*i.e.*, the cell was not otherwise occupied but lacked an egg). We avoided patches at the edge of the brood area. If a patch included an occasional cell with a newly eclosed larva, it was counted as ‘laid’ since it was likely that the eggs were on the verge of hatching. This method does not distinguish between eggs that were laid and then cannibalized by workers; however, since all colonies were fed supplemental protein, egg cannibalization should be linked to developmental deficiencies or failure to hatch rather than nutritional stress, which is a desirable feature to which our method should be sensitive. We repeated this procedure 2 weeks after the queens were exposed to pesticide treatments in order to calculate a change in laying pattern (the ratio of the fraction of cells laid post-stress relative to pre-stress).

On the day that we evaluated laying pattern, we caged queens with five attendants and candy, then transported to the laboratory where a colleague not otherwise involved in the study briefly anesthetized the queens with carbon dioxide, weighed them on an analytical balance, and randomized them into three treatment groups (untreated, 2 μl acetone, and 2 μl cocktail dissolved in acetone), keeping the lead experimenter blind to their assignments. The cocktail treatment was dosed at a hazard quotient (HQ) of 3,500, which is close to the median HQ that has been previously associated with ‘queen events’ [10]. This pesticide mixture consisted of the nine most abundant compounds found in the wax of commercial colonies, mixed and adjusted by serial dilution in the same relative proportions as reported by Traynor *et al*. [10]. This is the same mixture as was reported in McAfee *et al*. [42], but applied at a higher dose (**Table 3**). Treatments were administered topically to the queen’s thorax, then queens were returned to their cages with the workers, transported back to the apiary, and re-introduced to their respective colonies, but remained caged for 2 days before release. 2 weeks post-stress, laying patterns were again evaluated, the queens were transported to the laboratory, anesthetized, weighed, and sacrificed for dissection. Spermathecae were removed from the abdomen with forceps and blotted dry on a Kimwipe, then clean forceps were used to gently remove the tracheal net surrounding the spermatheca. The spermatheca was then lysed in an Ependorf tube containing 100 μl Buffer D and spermathecal fluid proteins were extracted for mass spectrometry analysis exactly as previously described [50].

We collected newly emerged (callow) workers from each colony 4 weeks after the beginning of the experiment and 4 weeks after the queens were stressed. It takes an average of 21 days for worker eggs to develop into adults [3]; therefore, callow workers collected 4 weeks after the beginning of the experiment developed from eggs laid approximately 1 week after the experiment began (*i.e.*, before the queens were stressed). Likewise, callow workers collected 4 weeks after the queens were stressed developed from eggs laid one week post-stress. The 2-day caging period immediately following the queen pesticide exposures provided time for the stress response to manifest and provided a short brood break to guard against overlap between pre- and post-stress newly emerged bees due to potential variation in developmental times.

We collected 9-12 callow workers per colony per time point in order to calculate a change in average mass at emergence. Callow workers are easily recognizable due to their light grey color, soft bodies, and inability to fly. We recorded the number of workers and wet weights on an analytical balance.

### Proteomics analysis

Proteins were extracted, digested, and purified from spermathecal fluid exactly as previously described [49, 50]. Briefly, sperm cells were spun down from the Buffer D-diluted spermathecal fluid solution and soluble proteins in the supernatant were precipitated with acetone. The pellets were washed and resuspended in urea digestion buffer (6 M urea, 2 M thiourea, in 100 mM Tris, pH 8). The proteins were reduced, alkylated, then digested with 0.2 μg of Lys-C (3 h, room temperature) followed by 0.2 μg of trypsin (overnight, room temperature, solution diluted with 4 volumes of 50 mM ammonium bicarbonate). Peptides were desalted using in-house made C18 STAGE-tips, dried, suspended in Buffer A (0.1% formic acid, 2% acetonitrile), and quantified using a Nanodrop (280 nm absorbance). One μg of peptides were injected on a Thermo easy-nLC 1000 liquid chromatography system coupled to a Bruker Impact II mass spectrometer. Sample orders were randomized prior to loading, and instrument parameters were set exactly as previously described [42]. We followed the same procedure for analysis of the queen fat bodies, except that protein was extracted into 6 M guanidinium chloride (in 100 mM Tris, pH 8) using a Precellys homogenizer with ceramic beads. We digested approximately 25 μg of protein per sample using 0.5 μg of Lys-C and trypsin.

Raw mass spectrometry data were searched using MaxQuant (v 1.6.1.0) exactly as previously described [42]. We used the most recent honey bee canonical protein database available on NCBI (HAv3.1, downloaded November 18^th^, 2019) with honey bee pathogen sequences added. Protein and peptide identifications were filtered to 1% FDR based on the reverse hits approach. All specific search parameters are available within the mqpar.xml file included in our data repository (see Data Availability).

### Statistical analysis

We analyzed laying pattern ratio, queen mass ratio, and worker mass ratio data using linear models in R (v3.5.1) with queen treatment included as a fixed effect. Queen sperm viability and morphometric analyses were conducted using R (v3.6.0). Because of the disparity between the number of tests to be run and the relatively small number of queens per group (n=4), we used principal components to reduce the number of variables under consideration. Namely, we generated the first principal component of all the correlated morphometric variables and used this as a final measurement of queen size. Queen size was used as a covariate in the final analyses. Additionally, we would not expect that sperm count would change as a result of this treatment, as it was applied after mating and non-viable sperm do not appear to be destroyed in the spermatheca [51]. Thus, we also used total sperm as a covariate in a final model testing the effect of topical agrichemical treatment on sperm viability.

For spermathecal fluid proteomics data, protein intensities (‘LFQ intensity’ columns from the MaxQuant output) were first log2 transformed, then reverse hits, contaminants, protein groups only identified by site, and protein groups without at least three defined values per treatment group were removed. Differential expression analysis was performed using limma() (example code is provided, see Data Availability) and a Benjamini-Hochberg multiple hypothesis testing correction to 10% FDR. We analyzed the queen fat body proteomics data in the same way, except queen fat body proteins were filtered to include only those that were identified in at least 3 of the 4 biological replicates at each dose. We analyzed expression of individual candidate biomarkers (six proteins) in the spermathecal fluid data using a linear model. Heatmaps were generated using Perseus v1.6.1.1 (clustered via Euclidian distance, 300 clusters, 10 iterations).

## Data availability

All raw proteomics data and search results have been deposited to the MassIVE proteomics repository (massive.ucsd.edu; field experiment data: accession MSV000086862; dose-response experiment data: accession MSV000087091). Phenotypic data for the queen dose-response topical exposure experiment are available in **Supplementary Table S1**. Queen fat body protein expression data and metadata are in **Supplementary Table S2 and S3**, respectively. Data from the queen exposure field trial is available in **Supplementary Table S4** and worker mass data is in **Supplementary Table S5.** An example R code for the limma protein expression analysis of fat bodies is available as **Supplementary File S1**. R code for the dose-response exposure analysis and field trial data analysis are available as **Supplementary File S2 and S3**, respectively.

## Acknowledgements

Funding from Project Apis m., Genome Canada (264PRO), Genome BC, and the BC Ministry of Agriculture supported this work. AM’s salary was supported by a Natural Sciences and Engineering Research Council fellowship. JPM was partially funded by a Graduate Student Fellowship through the NC Agricultural Foundation, and the entire project was supported from a grant from the Foundation for Food and Agriculture Research (Grant #549053). Special thanks to Jennifer Keller, Jordan Tam, Rhonda Thygesen, and Abigail Chapman for assistance with field work, dissections, and maintaining researcher blindness during the field trial.

## Notes

### Competing Interest Statement

The authors have declared no competing interest.

